# seagull: lasso, group lasso and sparse-group lasso regularisation for linear regression models via proximal gradient descent

**DOI:** 10.1101/2020.02.13.947473

**Authors:** Jan Klosa, Noah Simon, Pål O. Westermark, Volkmar Liebscher, Dörte Wittenburg

## Abstract

Statistical analyses of biological problems in life sciences often lead to high-dimensional linear models. To solve the corresponding system of equations, penalisation approaches are often the methods of choice. They are especially useful in case of multicollinearity which appears if the number of explanatory variables exceeds the number of observations or for some biological reason. Then, the model goodness of fit is penalised by some suitable function of interest. Prominent examples are the lasso, group lasso and sparse-group lasso. Here, we offer a fast and numerically cheap implementation of these operators via proximal gradient descent. The grid search for the penalty parameter is realised by warm starts. The step size between consecutive iterations is determined with backtracking line search. Finally, the package produces complete regularisation paths.

**Availability and implementation:** *seagull* is an R package that is freely available on the Comprehensive R Archive Network (*CRAN*; https://CRAN.R-project.org/package=seagull; vignette included). The source code is available on https://github.com/jklosa/seagull.

## 1 Introduction

Linear regression is a widely used tool to explore the dependence between a response variable and explanatory variables. For instance, in genome wide association studies (GWAS), the explanatory variables are typically the counts of genetic variants along the genome that might affect a response. The response variable could contain records of a disease or (continuous) measures of a performance trait. Deciphering the effect of the genetic variants and therewith uncovering the genetic architecture is essential in precision medicine and for animal or plant breeding programs.

The high throughput of modern biotechnological procedures enables studying an extremely large amount of explanatory variables (*p*). However, this often goes along with relatively few observations (*n*; *p* ≫ *n*), making the estimation of effects a challenge. Especially in the presence of multicollinearity, penalisation methods have proved to be useful for estimating effects. Famous examples are the *Tikhonov*, the *elastic net* (Zou and Hastie, 2005) and the *lasso* (Least Absolute Shrinkage and Selection Operator; Tibshirani, 1996) regularisation. In case only a very small fraction of variables has non-zero effect, those methods are advantageous that perform variable selection, such as lasso. However, in situations where the underling nature of effects is unknown, lasso might be prejudicial as it detects only the strongest signals. Then, causal relationships might be overlooked in GWAS (Waldmann *et al.*, 2013). To simultaneously detect non-zero effects and account for the relatedness of variables, the lasso has been modified and enhanced to the group lasso (Yuan and Lin, 2006), the sparse-group lasso (Simon *et al.*, 2013) and the ‘‘Integrative LASSO with Penalty Factors’’ (IPF-lasso, Boulesteix *et al.*, 2017). These particular modifications of the lasso assume an underlying group structure within the explanatory variables. A group structure is likely to appear in genomic data due to, for example, linkage and linkage disequilibrium. The R package presented here contains implementations of the lasso variants mentioned above focusing on precision of parameter estimation and computational efficiency.

## 2 Features

The R package *seagull* offers regularisation paths for optimisation problems of the general form:

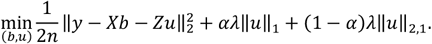

This is also known as the sparse-group lasso (Simon *et al.*, 2013). The first term expresses the “goodness of fit”. The second and third term are penalties, both of which are multiplied with the penalty parameter *λ* > 0. The vector *y* contains *n* observations of the response variable. The vectors *b* and *u* represent non-penalised and penalised effects, respectively; *X* and *Z* are the corresponding design matrices. Moreover, *α* ∈ [0, 1] is the mixing parameter for the penalties.

From a frequentist’s perspective, the non-penalised effects *b* and the penalised effects *u* are often referred to as *fixed* and *random effects*. Assuming a normal distribution for all occurring random variables, the above optimisation approach is equivalent to the maximisation of the penalised log-likelihood.

The two penalty terms are convexly linked via *α*. In the two limiting cases of *α* = 1 and *α* = 0, the respective resulting objective function is the lasso (Tibshirani, 1996) and the group lasso (Yuan and Lin, 2006). However, if *α* is chosen to be less than 1, it is assumed that the explanatory variables have an underlying group/cluster structure (with non-overlapping groups). The determination of such groups needs to be performed prior to the call of *seagull*, for instance, by applying a suitable cluster algorithm to the explanatory variables or by grouping them according to the source of measurement (RNA expression, SNP genotypes, etc.). Referring to this structure, the entries of *u* can be separated into the corresponding groups, say *u*^(*l*)^ for group *l*. Hence, the definition of the second penalty becomes:

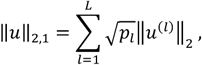

where *L* is the total number of groups and *p*_l_ is the size of group *l*. In our R package *seagull*, the following penalty operators are implemented: lasso, group lasso and sparse-group lasso. We show in the vignette of the package how the optimisation problem can be extended to consider weights for each explanatory variable and group. This option can be used for any reason but it also enables the user to apply the strategy of IPF-lasso.

## 3 Implementation

Both penalties are multiplied with the penalty parameter *λ* > 0; *λ* reflects the strength of the penalisation. Our package provides the opportunity to calculate a maximal value for *λ* and to perform a grid search by gradually decreasing this value. To efficiently accelerate this grid search, we implemented *warm starts*. This means, the solution for the current value of *λ* is used as starting point for the subsequent value of *λ*. Eventually, *seagull* provides a sequence of penalty parameters and calculates the corresponding path of solutions.

Furthermore, the above optimisation problem is solved via *proximal gradient descent* (PGD; e.g. Parikh and Boyd, 2014). PGD is an extension of gradient descent for optimisation problems which contain non-smooth parts, i.e. problems where the gradient is not available for the entire objective function. As PGD is an iterative algorithm, a proper step size between consecutive iterations is crucial for convergence. Determining such a step size is realised with *backtracking line search*. All implemented algorithms are based on the R package *Rcpp 1.0.3* (Eddelbuettel *et al.*, 2019).

## 4 Case study

We analysed the blood DNA methylation profiles at about 1.9 million CpG sites and its association with chronological age in a stock of mice (*n* = 141). The data set is publicly available and described in detail in Petkovich *et al.* (2017). Such data are used to build regression models termed epigenetic clocks, which enable biological age to be predicted from DNA methylation status. Standard approaches employ elastic net regression, which performs well but typically results in only ∼100 CpG sites with non-zero effect, limiting the potential for their genome-wide annotation and interpretation (Bell *et al.*, 2019). We split the data set into training (*n* = 75) and validation (*n* = 66) data, where all age classes appeared almost equally in both sets, and applied the sparse-group lasso variant of *seagull 1.0.5*. R scripts for the processing of the data are available in the supplementary material.

Fig. 1A shows the model fit based on regression coefficients which led to the minimum mean squared error of chronological age in the validation set. The correlation between the chronological and the predicted age (“Methylation age”) was 95.8%, and 5095 non-zero effects were identified. Hence using only the identified fraction of CpG sites enabled a precise prediction of age. As an option for regulating the sparsity, increasing the convergence parameter of *seagull* by two magnitudes (10^−6^ to 10^−4^) increased the number of non-zero effects by one magnitude. Furthermore, we compared the outcome of *seagull* to that of the established R package *SGL 1.3* (Simon *et al.*, 2019). Its implementation is based on accelerated generalized gradient descent. Though the implemented convergence criteria differed between both packages, results were similar. The correlation between regression coefficients leading to the minimum mean squared error was 99.5% (Fig. 1B). The number of non-zero effects obtained with *SGL* was 8822. In contrast to *SGL, seagull* computed the solution in a fraction of the time (*seagull*: ∼2 hours; *SGL*: ∼45 hours). In summary, *seagull* is a convenient envelope of lasso variants. Despite the similarities with *SGL*, only *seagull* offers the opportunity to incorporate weights for each penalised variable which enables further variants of the lasso.

**Figure 1.**
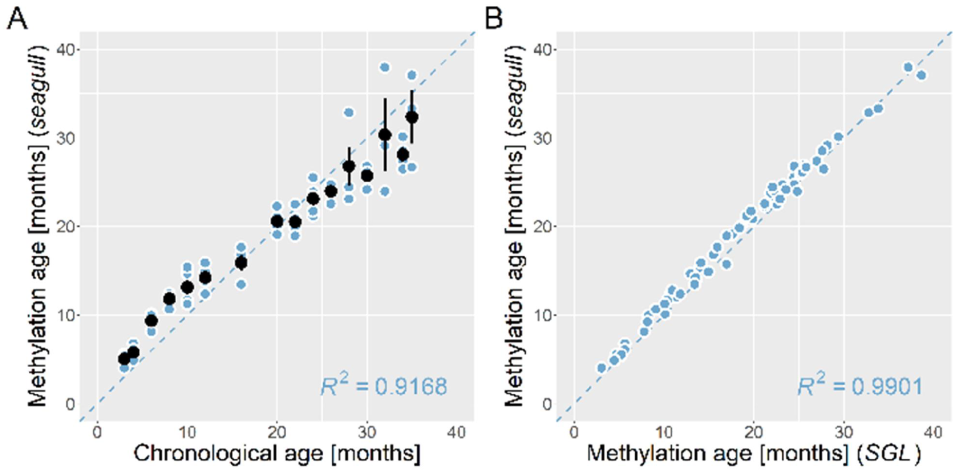
(A) Relationship between observed (chronological) and predicted (methylation) age. Each blue dot represents a sample in each class of observed chronological age (3mos, 4mos, etc.). Mean methylation age and error bars are displayed in black for each class of age. **(B) Methylation age obtained with seagull vs. SGL.** Blue dots represent samples; the dashed line is a regression line with slope 1.

